# Beak wiping stereotypies are correlated with neophobia and lack of enrichment in captive house sparrows (*Passer domesticus*)

**DOI:** 10.1101/2025.10.10.681718

**Authors:** Danna F. Masri, William J. Frazier, Melanie G. Kimball, Christine R. Lattin

## Abstract

The existence of behavioural syndromes, or suites of correlated behaviours, means that animals may not be able to act optimally in every situation, as they can constrain plasticity. Therefore, understanding links between different behaviours is critical for understanding why animals sometimes fail to respond appropriately to environmental challenges. In this study, we assessed whether beak wiping, a stereotyped anxiety-linked behaviour where birds wipe their beaks on a perch in a “windshield wiper” motion, was correlated with another anxiety-linked behavior, neophobia towards novel objects presented with food, in captive house sparrows (*Passer domesticus*). We predicted that more neophobic sparrows would also exhibit more beak-wiping stereotypies. We analyzed 1 h long control videos (when sparrows were presented with a normal food dish only; n=54) from three previous neophobia studies to assess beak wiping frequency, mean beak wiping bout duration, and total bout duration. Sparrows’ reluctance to feed in the presence of novel objects was significantly correlated with the mean duration of beak wiping bouts during control trials. We also found that simple enrichment (rubber perches, manzanita branch perches, and/or artificial pine branches) decreased both the frequency and duration of beak wiping. These findings suggest that neophobia and stereotypies may arise due to similar neuroendocrine mechanisms as part of a “high anxiety” behavioural syndrome. This work also highlights the importance of providing species-appropriate environmental enrichment to decrease the prevalence of stereotypic behaviours in captive songbirds.

## 1. Introduction

Consistent individual differences in behavior – known as personalities, temperaments, behavioral syndromes, or coping styles – have been described in dozens of different animal species [1-5]. An animal’s personality affects many behaviors that impact its fitness, from finding food [6, 7] to coping with competition [8, 9]. The existence of behavioural syndromes means that individuals may not be able to act optimally in every situation, as they can act as a potential constraint upon behavioral plasticity [10]. Therefore, understanding links between different behaviours is critical for understanding why animals sometimes fail to respond appropriately to environmental challenges.

One potential challenge wild animals can face is being transferred from the wild to a captive environment for research, conservation, or education purposes. The transition to captivity is often highly stressful for wildlife, partly due to differences between natural environments and controlled lab or zoo conditions, including forced proximity to humans, novel foods, artificial lighting, and restricted movement [11]. Captivity stress in turn can cause physiological and behavioural changes, such as the development of stereotypies [12, 13]. Stereotypies are short, repetitive sets of behaviours that replace the comparatively longer and more diverse range of behaviours observed in wild animals [14]. For example, because flight is inhibited by small cages, somersaulting stereotypies (where birds perform an aerial backflip off their perches, land on the cage floor, and fly back up to repeat the loop) may develop in captive birds as a response to thwarted escape behaviours [15]. Alternatively, some stereotypies involve natural behaviours that become disassociated from their natural context, such as beak wiping stereotypies that can develop in captive house sparrows (*Passer domesticus*) [16]. Beak wiping is a natural avian behaviour used to clean the beak after feeding, but in captivity it can become disassociated from feeding and much more frequent.

Stereotypic behaviour of captive animals is of scientific interest for several reasons. First, the performance of stereotypy can be used as a proxy for animal welfare, where less stereotypic behaviour is often (but not always) associated with better welfare [17, 18]. Providing environmental enrichment that allows for greater expression of natural behaviours such as movement, hiding, and grooming can substantially reduce stereotypies [23]. For example, European starlings (*Sturnus vulgaris*) housed in larger and longer cages that allowed for more flight exhibited fewer stereotypies compared to starlings in smaller cages [24]. Stereotypies can also impose a threat to the external validity and replicability of research on captive animals; if stereotypy induces behavioural inhibition, then behaviour-based assessments such as open field tests, reactions to novelty, or extinction and reversal learning may be affected [19]. However, not all wild species housed in captivity develop stereotypies [25], and there is wide individual variation within species in the amount of stereotypic behaviours performed in captive environments [26-28]. For example, licking stereotypies varied from 19% to 35% of observations among four captive northern giraffes (*Giraffa camelopardalis*) housed in an Indian zoo [29]. This across- and within-species variation suggests stereotypies may be related to variation in animals’ underlying neurobiology or physiology.

In this project, we investigated the connection between neophobia (novelty avoidance) and beak wiping stereotypies in captive house sparrows. We predicted that neophobia and beak wiping stereotypies might be linked as part of a high-anxiety behavioural syndrome resulting from shared neuroendocrine mechanisms [30]. For example, there is wide individual variation in glucocorticoid production in house sparrows in the wild and in captivity, and experimental evidence has linked glucocorticoids both to neophobia and to the development of beak wiping stereotypies in this species [31, 32]. We watched control videos from three previous studies of captive wild house sparrows, quantified the frequency, total duration, and mean duration of beak wiping bouts, and assessed whether any of these measures were correlated with a previously assessed measure of neophobia in the same birds, average latency to feed in the presence of novel objects [33-35]. Analyzing control videos (where the familiar food dish was replaced without novel objects) allowed us to assess beak wiping in a context separate from neophobia. Meta-analysis approaches find that most correlated behaviours show small effect sizes [36, 37], which suggests that many studies do not have sufficient sample size to reveal behavioural syndromes. Combining data from multiple studies provided us with a relatively robust sample size (n=54 birds) to look for evidence of a behavioural syndrome.

Our overall hypothesis was that neophobia and stereotypy are behaviours with similar underlying neuroendocrine mechanisms; therefore, we predicted that object neophobia and beak wiping would be positively correlated. Because housing conditions in these three studies varied slightly, this also gave us the opportunity to determine the effects of captive enrichment on stereotypies in wild-caught songbirds. We predicted that species-appropriate environmental enrichment (rubber perches, manzanita branch perches, and/or artificial pine branches) would decrease sparrow beak wiping bouts by allowing for more naturalistic behaviours (e.g., hiding behind artificial pine branches), thereby decreasing frustration and attenuating the need for stereotypy [24].

## 2. Methodology

### 2.1 Sparrow capture and enrichment

Adult house sparrows (n=54, 39 males and 15 females) used in this study were caught in three cohorts in East Baton Rouge Parish using mist nets; cohort capture dates were: A) June-July 2019 (n=22) [33], B) April-July 2022 (n=12) [35], and C) February 2022-April 2023 (n=20) [34]. Sparrows in Cohort A were part of a study examining the effects of exposure to novel objects or control conditions on neuronal activity; control videos used in the present study were from novel object, novel food, or novel object habituation trials [33]. Sparrows in Cohort B were part of a study aiming to understand the function of the avian hippocampus in neophobia. Control videos used for the present study were from novel object testing prior to the administration of any experimental treatments [35]. Finally, sparrows in Cohort C were part of a study investigating the effect of conspecific alarm calls on neophobia; we used control videos from the initial week of novel object trials before sparrows were exposed to calls [34].

Birds in all cohorts were allowed to habituate to captivity for at least three weeks before behaviour trials began. All sparrows were collected under Louisiana Scientific Research and Collecting Permits, and the Louisiana State University Institutional Animal Care and Use Committee approved all protocols (96-2018, 10-2021, 56-2022, 92-2023). To minimize the effect of social learning on neophobia [38], sparrows were singly housed in cages measuring 56 cm x 45 cm x 33 cm in the Louisiana State University vivarium. All sparrows had unlimited access to grit, mixed seeds, a vitamin-rich food supplement (Mazuri small songbird diet), and water; however, cohorts differed in the amount of environmental enrichment provided. For Cohort A, cages contained a plastic perch extending from the front to the back of the cage. For Cohort B, cages had the same plastic perch, a perch made of rubber tubing positioned diagonally, and a manzanita branch perch. Finally, the cages of Cohort C included the plastic and rubber tubing perches, plus an artificial pine branch hanging in the back of the cage (added as a possible hiding spot for sparrows) (**Fig 1**.) Note that prior to neophobia trials all sparrows also had a dish of sand for dustbathing, but these were removed during neophobia trials because they often contained food.

**Figure 1.**
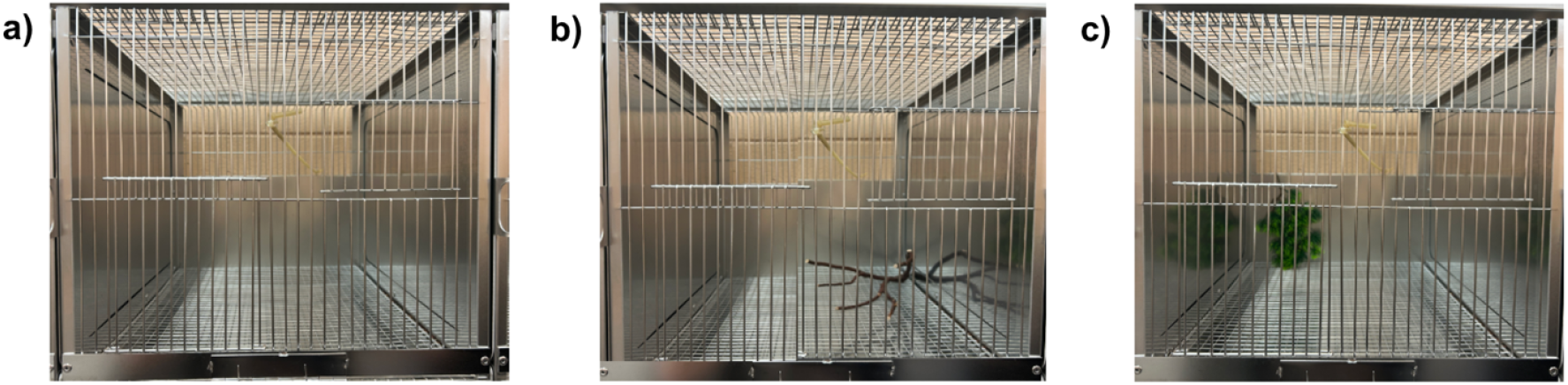
Environmental enrichment provided to house sparrows in: a) Cohort A (n=22); b) Cohort B (n=12); and c) Cohort C (n=20). All cohorts had a central longitudinal plastic perch. Cohort A received no additional enrichment during behaviour trials. Cohort B also received a manzanita branch perch and a diagonally-spanning rubber perch; Cohort C received an artificial pine branch and a diagonally-spanning rubber perch.

### 2.2 Behaviour analysis and quantification

In all three cohorts, sparrows were randomly assigned to different novel objects presented with food (n=3) or food alone as a control (n=2) during a week of novel object trials. For the three studies from which videos were pulled, a random number generator was used to select one of the control videos per sparrow to analyze for beak-wiping behaviours (n=54, 1 control video per sparrow). Each sparrow’s neophobia was quantified as previously published [33]; briefly, this was defined as an individual’s average latency to feed from the food dish when a novel object was present during three 1 h trials. To ensure that birds were motivated to feed during trials, food was removed overnight, and trials were conducted the following morning 30 min after lights on. To quantify neophobia, observers watched videos and recorded the bird’s first time eating from the food dish. If the bird did not feed, it was assigned a maximum time of 1 h. After the 1 h trials were complete, video recordings were stopped, all objects removed, and the normal food dish was present for the rest of the day.

Beak wiping, which we defined as a bird repeatedly wiping its beak back and forth in a “windshield wiper” motion, was used as a measure of stereotypy; each single event (dubbed a “bout”) was separated by 2 s [16]. Beak wiping stereotypy was recorded using BORIS software [39]. Two observers watched 1 h control videos and pressed a key to signify the beginning of a bout. When the bout ended, a key was pressed to indicate the end of the bout. This gave us three measures of stereotypy related to beak wiping: mean bout duration, total bout duration, and beak wiping frequency. Mean bout duration represents the average amount of time a bird spent in a single beak-wiping bout. Total bout duration is the cumulative time of all beak-wiping bouts recorded over a 1 h period. Finally, beak wiping frequency refers to the total number of beak-wiping bouts (each separated by at least 2 s) observed within an hour. Past research on beak wiping in house sparrows [16, 32] only counted numbers of bouts; however, because bouts can differ widely in length (e.g., if a sparrow performs two sets of wipes vs. twenty sets of wipes) and a key part of stereotypy is its repetitive nature, we decided to quantify beak wiping using multiple measures to determine if any aspect of this stereotypical behaviour was more clearly linked to neophobia or enrichment availability.

### 2.3 Statistical analyses

Analyses were run in in JMP Student Edition 18.2.2 (SAS Institute). We used three different sets of linear mixed models (one for each measure of beak wiping) to analyze the relationship between beak wiping and average latency to feed from a food bowl when novel objects were present. Initially, we tested whether the different types of enrichment found in Cohorts B and C (rubber tubing and a manzanita branch, and rubber tubing and an artificial pine branch, respectively) mattered for beak wiping behaviour, but because these two groups did not differ from each other (data not shown) they were combined as a “high enrichment” group (relative to the “low enrichment” Cohort A). For each model, we included fixed model effects of sparrow sex (male vs. female; determined by clear plumage features [40]), high enrichment (Cohort A = no, Cohorts B and C = yes), video quality (some videos were very dark or blurry and the two watchers made notes of this in their data sheets; poor quality vs. not poor quality), and average latency to feed in the presence of three novel objects (in s) and added video watcher as a random effect. We ran linear mixed models using Restricted Maximum Likelihood (REML) with unbounded variance components to prevent bias in estimation of fixed effects, and inspected residual plots to check that the homoscedasticity and linearity assumptions of linear models were met.

## 3. Results

Mean beak wiping bout duration (*F*_1,49_=4.48, p=0.040) was correlated with neophobia, while beak wiping frequency (*F*_1,1.6_=1.09, p=0.43) and total beak wiping duration (*F*_1,49_=0.86, p=0.36) were not (**Fig. 2, Table 1)**. Thus, sparrows that took longer to feed in the presence of novel objects also tended to have longer beak wiping bouts. Of the other fixed effects we examined in relation to beak wiping stereotypies (i.e., sparrow sex, presence of enrichment, and video quality), only environmental enrichment was significantly associated with all three measures of beak wiping **(Fig. 3, Table 1)**, where birds with high enrichment (i.e., Cohorts B and C) spent less total time beak wiping, beak wiped less frequently, and had shorter beak wiping bouts compared to birds in Cohort A. Video quality and sex were also correlated with mean wipe duration, where observers watching darker or lower-quality videos recorded lower mean bout times (*F*_1,49_=7.35, p=0.0092, **Fig. 4, Table 1**) and males had longer beak wiping bouts than females (*F*_1,49_=4.72, p=0.035).

**Table 1.** Fixed effect results from linear mixed models assessing the effects of captive house sparrow sex, high enrichment, video quality, and average latency to feed in the presence of three different novel objects (neophobia) compared to three different measures of beak wiping stereotypies assessed during a 1 h control video. Video watcher was included as a random effect in the model, and allowing for unbounded variance components affected degrees of freedom in some models. Significant effects are noted with a *.

|  |  | DF | DFDen | F Ratio | Prob > F |
| --- | --- | --- | --- | --- | --- |
| Mean wipe duration | Sex | 1 | 49 | 4.72 | 0.035 |
|  | Increased enrichment | 1 | 48 | 62.08 | <0.0001* |
|  | Video quality | 1 | 49 | 7.35 | 0.0092* |
|  | Average latency to feed in the presence of novel objects (neophobia) | 1 | 49 | 4.48 | 0.040* |
| Beak wiping frequency | Sex | 1 | 22 | 0.021 | 0.89 |
|  | Increased enrichment | 1 | 49 | 4.33 | 0.043* |
|  | Video quality | 1 | 33 | 0.85 | 0.36 |
|  | Average latency to feed in the presence of novel objects (neophobia) | 1 | 1.6 | 1.09 | 0.43 |
| Total wipe duration | Sex | 1 | 49 | 0.55 | 0.46 |
|  | Increased enrichment | 1 | 49 | 20.55 | <0.0001* |
|  | Video quality | 1 | 49 | 2.17 | 0.15 |
|  | Average latency to feed in the presence of novel objects (neophobia) | 1 | 49 | 0.86 | 0.36 |

**Figure 2.**
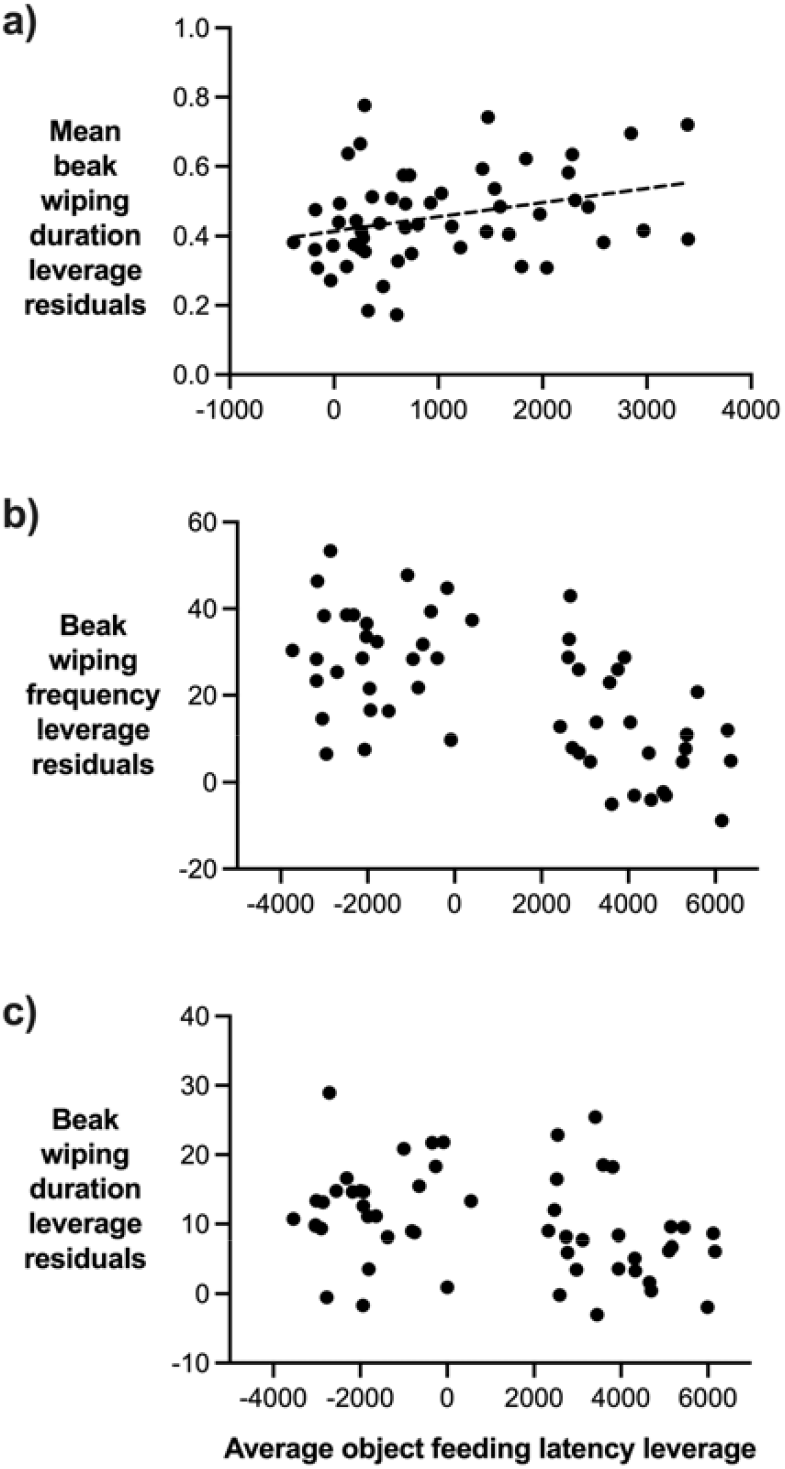
Leverage plots of a) mean beak wiping bout duration, b) beak wiping frequency, and c) total beak wiping duration in a 1 h control video (when the normal food dish was replaced) relative to house sparrows’ average latency to feed in the presence of novel objects (n=3 trials/bird, n=54 sparrows total). Mean beak wiping bout duration (a) was positively correlated with average object feeding latency, while beak wiping frequency and total bout duration were not related to neophobia. The y-axis is plotted as standardized leverage residuals of each data point, with each dot representing one sparrow.

**Figure 3.**
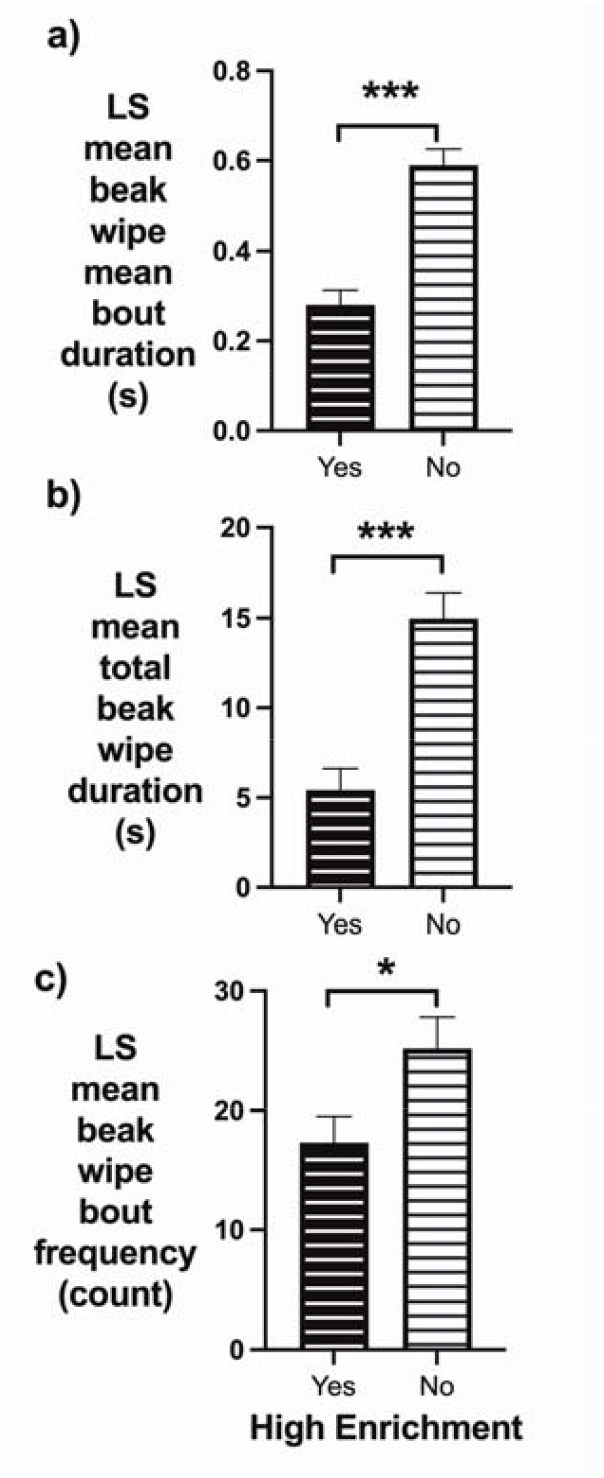
Least square means of the fixed effect variables of: a) increased enrichment on mean beak wipe duration; b) increased enrichment on total beak wipe duration; c) increased enrichment on beak wiping frequency in captive house sparrows during 1 h control videos (n=54). The presence of more enrichment (rubber perches and either manzanita branches or artificial pine branches) in Cohorts B and C significantly reduced all measures of beak wiping stereotypies compared to birds in Cohort A. * denotes p ≤ 0.05 and *** denotes p ≤ 0.0001.

**Figure 4.**
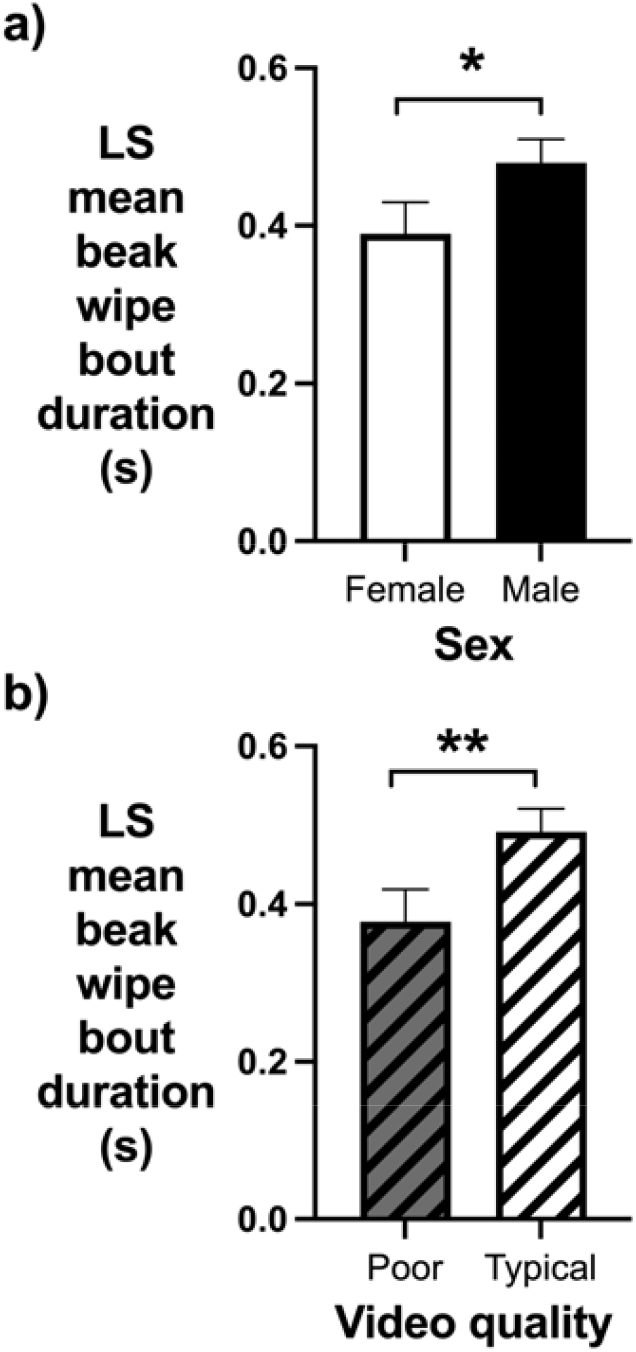
Least square means of the fixed effect variables of: a) sex and b) video quality on mean beak wiping duration in captive house sparrows during 1 h control videos (n=54). Both female sparrows and dark or blurry videos led to shorter mean beak wiping bouts. * denotes p ≤ 0.05 and ** denotes p ≤ 0.01.

## 4. Discussion

As predicted, neophobia and beak wiping stereotypy were correlated, though only for one of the three measures of beak wiping we tested, mean bout duration. Although little previous research has investigated within-individual associations between stereotypies and neophobia, studies examining links between stereotypy and a preference for or attraction to novel stimuli (neophilia) have found mixed results [26, 27, 41]. (Note that the opposite of neophobia is probably not neophilia, but indifference towards novelty; previous work suggests that neophobia and neophilia are distinct axes of behavior [42, 43].) Glucocorticoid production has been linked to both neophobia and stereotypy development in captive house sparrows [31, 32]; therefore, these two behaviours may be linked in a “high anxiety” behavioural syndrome via this shared hormonal mechanism, where, perhaps, sparrows with increased circulating levels of glucocorticoids or increased glucocorticoid receptor density might be more likely to be neophobic and have longer beak wiping bouts.

Another shared neurobiological mechanism between neophobia and stereotypies could involve the brain’s dopamine system, possibly with a permissive role for glucocorticoid hormones in dopamine secretion [32, 44]. When stimulated by amphetamine, dopamine receptors are activated and stereotypic behaviours induced, which can be intensified by high levels of corticosterone binding to glucocorticoid receptors [45]. Another study in captive house sparrows showed that giving the mixed dopamine D2/3 receptor agonist quinpirole resulted in a large and rapid reduction in beak wiping, showing a clear causal role for dopamine in this behaviour [32]. Further research is needed to test the role of individual variation in glucocorticoid and dopamine physiology in mediating a possible high-anxiety behavioural syndrome in house sparrows. A better understanding of the neurobiology of stereotypy in captive wild animals may help clarify the neurobiological basis for stereotypy in human mental health disorders such as obsessive-compulsive disorder (OCD) and autism [20-22].

As mentioned, only mean bout duration, and not beak wiping frequency or total bout duration, was significantly correlated with neophobia. This suggests that different measures of stereotypy may be more or less representative of an individual’s “average” stereotypy and likely to be linked to other behaviours in a syndrome. Also, behavioural syndrome research often emphasizes the importance of observing the same behaviour across different contexts to test whether a syndrome exists [46]. Because beak wiping observations took place exclusively during neutral (control) feeding contexts, follow-up studies could examine whether beak-wiping behaviours in more anxiety-inducing contexts—such as during novel food or object trials—are correlated with neophobia. Additionally, future studies may wish to use additional lighting in the bird room to minimize the impact of video quality on bout durations, as videos rated “poor quality” by watchers (these were usually a bit dark) consistently yielded shorter mean beak wiping bouts (note that an alternative explanation for these data is that birds in darker cages felt calmer, and performed less beak wiping as a result).

Finally, increased enrichment successfully reduced all three measures of stereotypy. Whether birds received rubber tubing and a manzanita branch (Cohort B) or rubber tubing and an artificial pine branch (Cohort C) did not matter: the presence of any additional level of environmental enrichment over that found in Cohort A significantly reduced beak wiping frequency, mean bout duration, and total bout duration. Overall, this study highlights the need for researchers to provide access to species-appropriate enrichment to maximize the wellbeing of songbirds and other research animals [24], while still being careful not to expose animals to novel objects, foods, or other stimuli that might induce neophobia and a stress response [47-49]. Appropriate environmental conditions are essential not only to ensure animal welfare in captivity, but also to elicit meaningful and generalizable responses to captive behaviour assays [19].

## Supporting information

Research data

## Acknowledgements

The authors thank animal caretakers and veterinary staff in the LSU Division of Laboratory Animal Research for sparrow husbandry and care, landowners who provided property access for capturing sparrows, and Juhee Haam and David Young for providing feedback on an earlier version of this manuscript.

## Data availability

Raw data are available as a supplementary .csv file.

## References

[1] B.E. Carlson, S.J. Tetzlaff, C. Rutz, Long□term behavioral repeatability in wild adult and captive juvenile turtles (Terrapene carolina): Implications for personality development, Ethology 126(6) (2020) 668–678.

[2] T.O. Cornwell, I.D. McCarthy, P.A. Biro, Integration of physiology, behaviour and life history traits: personality and pace of life in a marine gastropod, Animal Behaviour 163 (2020) 155–162.

[3] T.S.O. Costa, S.L.G. Nogueira-Filho, K.M. De Vleeschouwer, L.C. Oliveira, M.B.C. de Sousa, M. Mendl, L.S. Catenacci, S.S.C. Nogueira, Individual behavioral differences and health of golden-headed lion tamarins (Leontopithecus chrysomelas), American Journal of Primatology 82(5) (2020) e23118.

[4] A.M. Fisher, G.I. Holwell, T.A.R. Price, Behavioural correlations and aggression in praying mantids, Behavioral Ecology and Sociobiology 74(5) (2020).

[5] H.R. Thomson, S.D. Lamb, A.A. Besson, S.L. Johnson, Long-term repeatability of behaviours in zebrafish (Danio rerio), Ethology n/a(n/a) (2020).

[6] L.M. Aplin, D.R. Farine, R.P. Mann, B.C. Sheldon, Individual-level personality influences social foraging and collective behaviour in wild birds, Proceedings of the Royal Society B-Biological Sciences 281(1789) (2014) 20141016.

[7] U.A. Bergvall, A. Schäpers, P. Kjellander, A. Weiss, Personality and foraging decisions in fallow deer, Dama dama, Animal Behaviour 81(1) (2011) 101–112.

[8] D. Bierbach, C. Sommer-Trembo, J. Hanisch, M. Wolf, M. Plath, Personality affects mate choice: bolder males show stronger audience effects under high competition, Behavioral Ecology 26(5) (2015) 1314–1325.

[9] L.C. Garnham, S.A. Porthén, S. Child, S. Forslind, H. Løvlie, The role of personality, cognition, and affective state in same-sex contests in the red junglefowl, Behavioral Ecology and Sociobiology 73(11) (2019).

[10] A. Sih, A. Bell, J.C. Johnson, Behavioral syndromes: an ecological and evolutionary overview, Trends in Ecology and Evolution 19(7) (2004) 372–8.

[11] K. Morgan, C. Tromborg, Sources of Stress in Captivity, Applied Animal Behaviour Science 102 (2007) 262–302.

[12] G.J. Mason, Stereotypies: a critival review, Animal Behavior 41(6) (1991) 1015–1037.

[13] R.M. Calisi, G.E. Bentley, Lab and field experiments: are they the same animal?, Horm Behav 56(1) (2009) 1–10.

[14] D. Eilam, R. Zor, H. Szechtman, H. Hermesh, Rituals, stereotypy and compulsive behavior in animals and humans, Neuroscience & Biobehavioral Reviews 30(4) (2006) 456–71.

[15] G. Feenders, M. Bateson, The development of stereotypic behavior in caged European starlings, Sturnus vulgaris, Developmental Psychobiology 54(8) (2012) 773–784.

[16] C.R. Lattin, A.V. Pechenenko, R.E. Carson, Experimentally reducing corticosterone mitigates rapid captivity effects on behavior, but not body composition, in a wild bird, Hormones and Behavior 89 (2017) 121–129.

[17] D.L. Wells, Sensory stimulation as environmental enrichment for captive animals: A review, Applied Animal Behaviour Science 118(1-2) (2009) 1–11.

[18] G.J. Mason, N.R. Latham, Can’t stop, won’t stop: is stereotypy a reliable animal welfare indicator?, Animal Welfare 13(S1) (2004) S57–S69.

[19] J.P. Garner, G.J. Mason, Evidence for a relationship between cage stereotypies and behavioural disinhibition in laboratory rodents, Behavioural Brain Research 136(1) (2002) 83–92.

[20] M. Low, Stereotypies and behavioural medicine: confusions in current thinking, Australian Veterinary Journal 81(4) (2003) 192–8.

[21] J.P. Garner, C.L. Meehan, J.A. Mench, Stereotypies in caged parrots, schizophrenia and autism: evidence for a common mechanism, Behavioural Brain Research 145(1-2) (2003) 125–34.

[22] S.D. Fam, Y.S. Tan, C. Waitt, Stereotypies in Captive Primates and the Use of Inositol: Lessons from Obsessive–Compulsive Disorder in Humans, International Journal of Primatology 33(4) (2012) 830–844.

[23] R. Swaisgood, D. Shepherdson, Environmental enrichment as a strategy for mitigating stereotypies in zoo animals: a literature review and meta-analysis, Stereotypic animal behaviour: fundamentals and applications to welfare (2006) 256–285.

[24] M. Bateson, G. Feenders, The use of passerine bird species in laboratory research: implications of basic biology for husbandry and welfare, ILAR Journal 51(4) (2010) 394–408.

[25] G.J. Mason, Species differences in responses to captivity: stress, welfare and the comparative method, Trends in ecology & evolution 25(12) (2010) 713–721.

[26] S. Goswami, P.C. Tyagi, P.K. Malik, S.J. Pandit, R.F. Kadivar, M. Fitzpatrick, S. Mondol, Effects of personality and rearing-history on the welfare of captive Asiatic lions (Panthera leo persica), PeerJ 8 (2020) e8425.

[27] S. Silber, S. Joshi, N. Pillay, Behavioural syndromes in stereotypic striped mice, Applied Animal Behaviour Science 212 (2019) 74–81.

[28] M.L. Fangmeier, A.L. Burns, V.A. Melfi, J. Meade, Foraging enrichment alleviates oral repetitive behaviors in captive red□tailed black cockatoos (Calyptorhynchus banksii), Zoo biology 39(1) (2020) 3–12.

[29] T.P. Kulkarni, Analysis of stereotypic behaviour and enhanced management in captive Northern Giraffe Giraffa camelopardalis housed at Zoological Garden Alipore, Kolkata, Journal of Threatened Taxa 12(4) (2020) 15426–15435.

[30] J.M. Koolhaas, S.M. Korte, S.F. De Boer, B.J. Van Der Vegta, C.G. Van Reene, H. Hopster, I.C. De Jong, M.A.W. Ruis, H.J. Blokhuis, Coping styles in animals: current status in behavior and stress-physiology, Neuroscience & Biobehavioral Reviews (1999).

[31] T.R. Kelly, K.I. Lynch, K.E. Couvillion, J.N. Gallagher, K.R. Stansberry, M.G. Kimball, C.R. Lattin, A transient reduction in circulating corticosterone reduces object neophobia in male house sparrows, Hormones and Behavior 137 (2022) 105094.

[32] C.R. Lattin, D.P. Merullo, L.V. Riters, R.E. Carson, In vivo imaging of D(2) receptors and corticosteroids predict behavioural responses to captivity stress in a wild bird, Scientific Reports 9(1) (2019) 10407.

[33] M.G. Kimball, K. Lynch, K.E. Couvillion, J.N. Gallagher, K.R. Stansberry, M.G. Kimball, C.R. Lattin, Novel objects alter immediate early gene expression globally for ZENK and regionally for c-Fos in neophobic and non-neophobic house sparrows, Behavioural Brain Research 428 (2022).

[34] M.G. Kimball, D.F. Masri, E.B. Gautreaux, K.R. Stansberry, T.R. Kelly, C.R. Lattin, Conspecific alarm calls prevent the attenuation of neophobia behavior in wild-caught house sparrows (Passer domesticus), Frontiers in Bird Science 3 (2024).

[35] M.G. Kimball, Investigating the Effects of Environmental Perturbations on House Sparrow Neurobiology and Behavior, (2024).

[36] L.Z. Garamszegi, G. Markó, G. Herczeg, A meta-analysis of correlated behaviours with implications for behavioural syndromes: mean effect size, publication bias, phylogenetic effects and the role of mediator variables, Evolutionary Ecology 26(5) (2012) 1213–1235.

[37] L.Z. Garamszegi, G. Markó, G. Herczeg, A meta-analysis of correlated behaviors with implications for behavioral syndromes: relationships between particular behavioral traits, Behavioral Ecology 24(5) (2013) 1068–1080.

[38] T.R. Kelly, M.G. Kimball, K.R. Stansberry, C.R. Lattin, No, you go first: phenotype and social context affect house sparrow neophobia, Biology Letters 16(9) (2020) 20200286.

[39] O. Friard, M. Gamba, BORIS: a free, versatile open□source event□logging software for video/audio coding and live observations, Methods in Ecology and Evolution 7(11) (2016) 1325–1330.

[40] P. Lowther, C. Cink, S. Billerman, House sparrow (Passer domesticus), version 1.0, Birds of the World 1 (2020).

[41] S.S. Vickery, G.J. Mason, Stereotypy and perseverative responding in caged bears: Further data and analyses, Applied Animal Behaviour Science 91(3-4) (2005) 247–260.

[42] C. Mettke-Hofmann, H. Winkler, B. Leisler, The Significance of Ecological Factors for Exploration and Neophobia in Parrots, Ethology 108(3) (2002) 249–272.

[43] M.G. Kimball, C.R. Lattin, The “Seven Deadly Sins” of Neophobia Experimental Design, Integr Comp Biol (2023).

[44] F.N. Madison, V.P. Bingman, T.V. Smulders, C.R. Lattin, A bird’s eye view of the hippocampus beyond space: Behavioral, neuroanatomical, and neuroendocrine perspectives, Hormones and Behavior 157 (2024) 105451.

[45] J.R. Pauly, S.F. Robinson, A.C. Collins, Chronic corticosterone administration enhances behavioral sensitization to amphetamine in mice, Brain research 620(2) (1993) 195–202.

[46] A. Sih, A.M. Bell, J.C. Johnson, R.E. Ziemba, Behavioral syndromes: an integrative overview, The Quarterly Review of Biology 79(3) (2004) 241–277.

[47] S. Richard, N. Wacrenier-Ceré, D. Hazard, H. Saint-Dizier, C. Arnould, J. Faure, Behavioural and endocrine fear responses in Japanese quail upon presentation of a novel object in the home cage, Behavioural Processes 77(3) (2008) 313–319.

[48] R.A. Fox, J.R. Millam, Novelty and individual differences influence neophobia in orange-winged Amazon parrots (Amazona amazonica), Applied Animal Behaviour Science 104(1-2) (2007) 107–115.

[49] T.M. Meade, E. Hutchinson, C. Krall, J. Watson, Use of an aquarium as a novel enrichment item for singly housed rhesus macaques (Macaca mulatta), J Am Assoc Lab Anim Sci 53(5) (2014) 472–7.

